# Disinfectant residuals in drinking water systems select for mycobacterial populations with intrinsic antimicrobial resistance

**DOI:** 10.1101/675561

**Authors:** Maria Sevillano, Zihan Dai, Szymon Calus, Quyen M Bautista-de los Santos, A. Murat Eren, Paul W.J.J. van der Wielen, Umer Z. Ijaz, Ameet J. Pinto

## Abstract

Antimicrobial resistance (AMR) in drinking water has received less attention than counterparts in the urban water cycle. While culture-based techniques or gene-centric PCR have been used to probe the impact of treatment approaches (e.g., disinfection) on AMR in drinking water, to our knowledge there is no systematic comparison of AMR traits between disinfected and disinfectant residual-free drinking water systems. We use metagenomics to assess the associations between disinfectant residuals and AMR prevalence and its host association in full-scale drinking water distribution systems (DWDSs). The differences in AMR profiles between DWDSs are associated with the presence or absence of disinfectant. Further, AMR genes and mechanisms enriched in disinfected systems are associated with drug classes primarily linked to nontuberculous mycobacteria (NTM). Finally, evaluation of metagenome assembled genomes (MAGs) of NTM indicates that they possess AMR genes conferring intrinsic resistance to key antibiotics, whereas such NTM genomes were not detected in disinfectant residual free DWDSs. Thus, disinfection may not only influence the AMR profiles of the drinking water microbiome but also select for NTM with intrinsic AMR.

## INTRODUCTION

Regulation compliant drinking water can contain low concentrations of chemicals and microorganisms originating from source water, treatment process, distribution systems, and premises plumbing^1,2^. While our understanding of chemical contaminants (e.g., disinfection by-products (DBP)^3^) and microbial populations^4–6^ in treated drinking water is increasing, the drinking water field has only recently started investigating emerging biological contaminants^7^ like antibiotic resistance bacteria (ARB) and antibiotic resistance genes (ARG). Misuse of antibiotics and anthropogenic release of bioactive compounds into the urban water cycle exacerbate the prevalence and persistence of antimicrobial resistance (AMR) and can make engineered systems for water treatment (i.e., wastewater and drinking water) critical points of AMR dissemination^8,9^. Drinking water can constitute a potential exposure route to microorganisms with intrinsic and acquired antibiotic resistance genes (i.e., resistome^10^), and it is particularly concerning if AMR traits are associated with waterborne pathogens^11^.

Drinking water disinfection is one of the most important public health advances towards elimination of waterborne diseases and remains the most widely used treatment strategy today for pathogen control^12^. Typically, drinking water systems (DWS) maintain a disinfectant residual, usually chlorine, in the drinking water distribution system (DWDS) to minimize microbial growth. However, its unintended impacts (e.g., DBP formation) have prompted the evaluation of alternative strategies to mitigate associated risks. These range from the use of alternate disinfectants such as chloramines, to advanced oxidation processes that avoid the use of disinfectants and sometimes eliminate the need for maintaining a disinfectant residual. The latter approach, i.e., absence of disinfectant residual, is practiced in some European countries (e.g., Denmark, Netherlands, Switzerland, and parts of Germany) for production and distribution of safe drinking water^13^. Among the range of treatment strategies to manage microbial growth in the DWDS, disinfection is likely to have a significant effect on AMR prevalence; either via direct^14–16^ or indirect (i.e., DBP mediated) selection^17^ of resistant microbial populations. For instance, sub-lethal chlorine concentrations are associated with upregulation of ARGs in pathogens^14,15^ and can promote conjugative transfer of ARGs within strains and across genera^16^. Culture dependent work has shown that the presence of chlorine^18^ and monochloramine^19^ can result in increased ARB prevalence in the DWDS. Similarly, Shi *et al.* showed an increase in the relative abundance of ARGs after chlorination with subsequent decrease at the tap^20^. Xu *et al.* showed that a terminal chlorination step in advanced ozone-biological activated carbon treatment enhanced ARG concentrations^21^. Others have argued that the change in resistance profiles are likely driven by changes in bacterial community composition during drinking water treatment rather than selection for the resistance traits themselves^22^.

One of the key challenges with contextualizing the importance of ARGs in DWS has been the inability to resolve host-association of detected ARGs. The presence of ARGs in pathogenic microorganisms or mobile genetic elements can present higher exposure risk, as compared to their presence in non-pathogenic and/or innocuous microbes^23,24^. Although qPCR is the gold standard for ARG detection and quantitation, it requires *a priori* knowledge of the types of ARGs that need to be quantified and it is unable to determine ARG host association. In contrast, high throughput sequencing (HTS) (e.g., metagenomics) can be used to identify a broader range of ARGs and draw insights about the functionality of detected ARGs and their host association. Multiple studies have shown high concordance between these two techniques, and show that metagenomic methods can indeed be useful for relative quantitation^25–29^. However, there are key caveats associated with the use of metagenomic data. Cost of sequencing can be significantly higher than targeted qPCR with the trade-off being the ability to capture larger number of genes. While qPCR can detect target genes at extremely low levels, metagenomics is limited to detecting medium-to-high abundance genes with detection of rare traits requiring extremely deep sequencing effort^30^. Further, metagenomic based ARG identification is impacted by the use of arbitrary thresholds for functional annotation^31–34^, requiring additional curation efforts to ensure the trait assignment is robust.

In this study, we used a metagenomic approach to compare the prevalence of ARGs and their hosts in DWS with different disinfection strategies. To do this, we analysed tap water samples from several locations within mainland UK and the Netherlands that either maintain (Dis) or lack disinfectant residual (NonDis) in the DWDS. Specifically, our goals were to (1) survey ARGs across a range of Dis and NonDis drinking water distribution systems, (2) assess their abundance and diversity in the context of presence/absence of disinfectant residual, (3) employ *de novo* assembly and genome binning to determine the host-association of high prevalence ARGs, and (4) relate the prevalence of ARG harbouring microbes and/or mobile genetic elements to the presence/absence and concentration of a disinfectant residual in the DWDS.

## METHODS

### Drinking water sampling and water quality analyses

A total of 39 samples were collected from 11 drinking water systems (DWS) including systems with a disinfectant residual (Dis), located in the UK (n=21), and without a disinfectant residual (NonDis), located in the Netherlands (n=18). The Dis samples were collected during summer 2015 and NonDis samples were collected in the summer of 2013. A detailed sampling protocol was described previously^35^. Briefly, tap water was sampled after flushing the faucet for 20 min, to minimize effects from premises plumbing. A constant flow rate was maintained, and the grab sample was either collected from the tap in sterile (by autoclaving) Nalgene containers, transported to the laboratory and immediately filtered; or filtered onsite using sterile equipment. Drinking water was filtered in triplicate through 0.2 µm Sterivex filters (SVGP01050 EMD Millipore, USA) using a peristaltic pump (Watson-Marlow 323S/D, UK) until the filter clogged or up to a 15 L volume for each filter. Water quality parameters measurements were performed as described previously^35^. This included on-site measurements of temperature, pH, conductivity, and dissolved oxygen using an Orion 5 Star Meter (Thermo Fisher Scientific, USA). Total chlorine was measured on site with EPA approved HACH kit on a DR 2800 VIS Spectrophotometer (Hach Lange, UK). Nitrogen species (ammonia, nitrite, and nitrate) were measured in the laboratory using standard methods 4500-NH_3_-F, 4500-NO_2_-B, and 4500-NO_3_-B respectively. Total Organic Carbon (TOC) was measured with a Shimadzu TOC-LCPH Analyzer (Shimadzu, Japan). Detailed measurements and descriptions of sampling sites can be found in supplementary materials (Table S1).

### DNA extraction and shotgun sequencing

Filter membranes were aseptically removed from the Sterivex cartridge and transferred to 2 ml Lysing Matrix E tubes (SKU 116914100, MP Biomedicals, USA) and DNA extraction and purification were performed in a Maxwell® 16 DNA extraction system (Promega) using the LEV DNA kit (AS1290, Promega, USA). Briefly, 300 µl of lysing buffer and 30 µl of Proteinase K were added to the Lysing Matrix E tubes containing the filter membrane, followed by incubation at 56°C for 20 min. Subsequently, 500 µl of chloroform:isoamyl alcohol (24:1, pH 8.0) was added to the tube and the tube was vortexed, followed by bead beating for 40 s at 6 m/s using a FastPrep 24 instrument (MP Biomedicals, USA), and centrifugation at 14,000g for 10 min. The aqueous phase of the supernatant was transferred to a 2 ml centrifuge tube and two more bead beating steps were performed by replacing the aqueous phase with fresh lysing buffer (75 µl and 50 µl, respectively) prior to each bead beating step and followed by centrifugation at 14,000g for 10 min. Subsequent DNA purification from the aqueous phase was carried out by Maxwell LEV DNA kit. The extracted DNA was quantified using a Qubit HS dsDNA kit (Q32854, Life Technologies, UK) with a Qubit 2.0 Fluorometer (Life Technologies, UK). Negative controls consisting of reagent blanks (no input material) and filter blanks (filter membranes from unused Sterivex filters) were processed identically as the samples for DNA extraction (n=8).

Libraries were prepared using the Nextera XT DNA Sample Preparation Kit (FC-131-1096, Illumina Inc.) according to the manufacturer’s protocol. DNA extracts from the reagent and filter blanks were spiked with genomic DNA from an even and an uneven mock community consisting of genomic DNA from 10 organisms (Table S2) and included in the library preparation and sequencing run. All samples were cleaned up with HighPrep PCR magnetic beads (AC-60050, MagBio Inc.) according to manufacturer’s instructions to remove very short fragments, evaluated for fragment size using High Sensitivity DNA Kit on Agilent Bioanalyzer (5067-4626, Agilent Inc.), and then quantified with qPCR according to Illumina guidelines. Libraries from all samples and spiked negative controls were normalized based on qPCR results and pooled in equimolar concentration. Finally, the pooled samples were quantified with Qubit HS dsDNA assay and concentrated using HighPrep PCR magnetic beads. The sequencing library was then subject to metagenomic sequencing on four lanes of Illumina HiSeq 2500 flow cell (paired end, dual indexing, 2×250 bp read length, Rapid Run Mode) at University of Liverpool Centre for Genomic Research (CGR, Liverpool, UK).

### Metagenomic data processing

Quality filtering was performed at the CGR on the raw FASTQ files by trimming reads to remove Illumina adapter sequences using Cutadapt^36^ version 1.2.1. The option -O 3 was used, so the 3’ end of any reads which match the adapter sequence for 3 bp or more were trimmed. The reads were further trimmed using Trimmomatic^37^ v0.35 with a minimum Phred score of Q20. Filtered paired end reads were interleaved and co-assembled for each drinking water system using MetaSpades^38^ v3.10.1, for a total of 11 co-assemblies (6 Dis and 5 NonDis). The resulting scaffolds were filtered by size selection, and only scaffolds 500 bp or longer were used for downstream analyses. Coverage information was obtained by mapping trimmed reads from each sample against scaffolds with bwa-mem^39^ v0.7.12and followed by using genomecov from bedtools^40^. Furthermore, contaminant analyses were incorporated into the processing workflow as described by Dai *et al.* (in preparation). Briefly, a scaffold was considered present in a sample if it was not detected in the negative controls or if its relative abundance in sample was greater than the control and coverage across the length of scaffold in the sample was more uniform than in the negative control. A detailed explanation of this approach is provided in Supplementary text 1. Using this approach, each scaffold was labelled as a “true” scaffold (i.e., present in the sample) or a “contaminant” scaffold. Open reading frames (ORFs) were identified using prodigal^41^ v2.6.3. The predicted ORFs were then searched against *rpoB* specific hmm profile (pf04563) from the Pfam-A database using hmmsearch (hmmer.org). The *rpoB* normalized coverage of each true scaffold was determined by dividing its coverage in each sample by the cumulative coverage of all scaffolds containing *rpoB* genes in that sample. Scaffold level coverage was used for normalization purposes to account for unequal mapping density of reads across scaffolds.

### Annotation of Antibiotic Resistance Genes

Predicted open reading frames (ORFs) were mapped against the Comprehensive Antibiotic Resistance Database^42^ (CARD, homologous model, V1.1.3) using DIAMOND^43^ v0.8.2.64 to identify antibiotic resistance ontologies (ARO) using an approach analogous to reciprocal-BLAST. Specifically, the predicted ORFs were aligned to the CARD and then the CARD was aligned against the ORFs in amino acid space with a minimum query coverage of 70% (i.e., 70% of the ORF length was aligned to the CARD reference sequence and subsequently 70% of the CARD reference sequence was aligned against the ORF sequence) using DIAMOND. This was followed by manual curation of sample vs CARD and CARD vs sample alignments to assign AROs to ORFs that demonstrated best match to the CARD reference sequence and consistent annotation using both alignment approaches. Following this curation, ARO tables were generated for three amino acid percent identity thresholds (i.e. 30%, 50%, and 70%) consisting of the *rpoB* normalized coverage of scaffolds identified as containing an ARO at each percent identity threshold. The three percent identity thresholds are referred to as “loose” (30%), “medium” (50%), and “stringent” (70%) sequence similarity thresholds. This approach was taken because some proteins can provide functional resistance to antibiotics even at low percent identity thresholds to known ARGs^31^. Indexing files from the CARD were used to categorize results by target drug class and mechanism of resistance, and any ARO annotated whose name string contained “mutant” was removed from the analysis to avoid inclusion of resistance conferred by mutations.

### Identifying host and mobile genetic element association of ARGs

All scaffolds were classified with Kaiju^44^ v1.4 using the ncbi-nr database (RefSeq Version 86) as reference database and followed by web-based BLASTn on NCBI-nt database to confirm Kaiju assigned taxonomic annotation for select important scaffolds. Second, all scaffolds containing AROs were aligned against a local database of genomes of WHO identified waterborne pathogens (Bacteria, Viruses, Protozoa and Helminths) downloaded from NCBI (RefSeq Version 86) using BLASTn with a minimum query coverage and percent identity thresholds of 90% (Supplementary text 2). This database also contained genomes from bacterial groups that have been previously identified as indicators of the antibiotic resistance status in the environment^8^. Scaffolds that contained AROs but did not show significant similarity to the genomes in pathogen database (90% identity and 90% query coverage) were further analyzed on read basis. Specifically, reads mapping to these scaffolds were extracted from corresponding BAM files and these reads were aligned against the pathogen reference database using BLASTn with sequence similarity threshold of minimum 95% identity and 95% query coverage. Finally, we also analyzed the ARO containing scaffolds to determine if they were associated with mobile genetic elements by determining if they were likely plasmid or viral scaffolds. To do this, the scaffolds were aligned against a local database consisting of plasmid sequences or viral sequences obtained from NCBI (RefSeq Version 86) with an identity and query coverage threshold of 97% and 70%, respectively. Further, the scaffolds containing AROs were analyzed for viral origin by (1) aligning against the IMG/VR database^45^ using BLAST with an identity and query coverage threshold of 97% and 70%, respectively, (2) using the k-mer based approach of VirFinder^46^ with default settings, and (3) using the IMG/VR protocol for viral detection using the virus discovery pipeline as outlined previously^47^.

### Genome binning, annotation, and phylogenomics

Metagenome assembled genomes (MAGs) were recovered by binning using the CONCOCT^48^ implementation in anvi’o^49^. Resulting bins were refined using RefineM^50^ and then manually curated in anvi’o and de-duplicated by dRep^51^. The quality of all refined bins was then assessed by CheckM^52^. Taxonomic assignment was performed based on GTDB-Tk^53^ v0.1.3. The web-based Resistance Gene Identifier (RGI) tool of the CARD was used to confirm that select bins carried identified ARGs. DNA sequences were queried to recover perfect or strict hits only (according to the CARD nomenclature) and the high quality/coverage option was selected. Phylogenomic trees were generated in anvi’o by first using the program ‘anvi-get-sequences-for-hmm-hits’ to recover individually-aligned and concatenated 48 single-copy ribosomal protein genes^54^ from five mycobacterial MAGs from this study and 35 complete mycobacterial reference genomes downloaded from NCBI (Table S3), and then using the program ‘anvi-gen-phylogenomic-tree’ that infers evolutionary associations between genomes using FastTree^55^ v2.1.7.

### Data analyses and statistics

Statistical analyses were conducted in R software^56^ and visualizations generated with ggplot package. Non parametric testing was performed with R base statistic packages using function wilcox.test(). Alpha diversity metrics were obtained with sepecnumber() and diversity() functions from vegan^57^ package 2.4-0. Beta diversity was evaluated with pairwise comparisons of *rpoB* normalized ARO abundances using Bray Curtis distances in metaMDS() function from vegan. Permutational ANOVA was performed using the perm.oneway.anova() function from wPerm package (https://cran.r-project.org/web/packages/wPerm/index.html). PERMANOVA tests were conducted using the adonis() function from vegan. A SIMPER analysis was perform in order to evaluate ARO contributions to the dissimilarity between system type using simper() function.

## RESULTS AND DISCUSSION

To investigate the prevalence of antibiotic resistance genes (ARGs) as a function of disinfection strategy (i.e., disinfectant residual present (Dis) vs disinfectant residual free (NonDis)) we generated 11 shotgun metagenomes from 39 samples that contained on average 7.39 ×10^6^ paired-end reads per sample after quality control (Supplementary Table S4) and an average metagenome assembly size of 434.4 Mbp. Further, gene annotations were inspected at different amino acid percent identity thresholds categorized as “loose” (percent identity ≥ 30%), “medium” (percent identity ≥ 50%), and “stringent” (percent identity ≥ 70%) to evaluate protein similarity and provide a comprehensive exploration of detected antimicrobial resistance (AMR) traits^58^.

### Antimicrobial resistance traits are prevalent in drinking water systems irrespective of presence/absence of disinfectant residual

We first searched all predicted gene sequences from open-reading frames in assembled scaffolds in the Comprehensive Antibiotic Resistance Database (CARD) with a query coverage threshold of 70% and sequence similarity threshold of 30%. These criteria resulted in 18256 hits to investigate the distribution of the antibiotic resistant ontology (ARO) in drinking water system (DWS) samples in Dis and NonDis groups (Figure 1A, Supplementary Table S5). We describe semi-quantitative differences between Dis and NonDis groups through normalizing the coverage of AROs within samples by the sample’s *rpoB* gene coverage. The *rpoB* gene, unlike the 16S rRNA gene, is a single copy gene that can be used as a molecular marker to estimate the bacteriological diversity of a sample and normalize the abundance of functional genes of interest^59–61^. While fewer scaffolds from Dis samples contained AROs (i.e., ∼39% of all significant hits) compared to the NonDis samples, the mean *rpoB* normalized coverage of these ARO containing scaffolds was significantly higher (p < 0.001) in the Dis (9.3±20×10^−2^) compared to the NonDis (1.6±4×10^−2^) samples. Of the 451 AROs detected in this study, a total of 322 and 397 AROs were present in Dis and NonDis samples, respectively. Further, a greater number of shared AROs between the two systems were more abundant in Dis samples (n=213) compared to NonDis samples (n=55) (Figure 1B). Dis systems have fewer exclusive AROs (n=54) compared to NonDis samples (n=129), with a large number of detected AROs shared between the two groups (n=268) (Figure 1C). A comparison of the mean *rpoB* normalized coverage of AROs between the two groups (i.e., Dis vs NonDis) that were present in more than 50% of the samples for each strategy, indicated that most shared AROs were more abundant in the Dis samples compared to the NonDis samples. (Figure 1D). Only three shared and frequent AROs (>50% detection frequency) were more abundant in NonDis as compared to Dis samples; these AROs included *vanHB, dfrE*, and *clbB*. The top three AROs that were significantly more abundant in Dis compared to NonDis systems were *vanRB, iri, oprN*. The most significant differentially abundant ARO in Dis samples was associated with multidrug efflux pump *sav1866*, which was detected in all samples analyzed and had a mean *rpoB* normalized coverage of 1.49±0.95 and 0.37±0.24 in Dis and NonDis samples, respectively.

**Figure 1:**
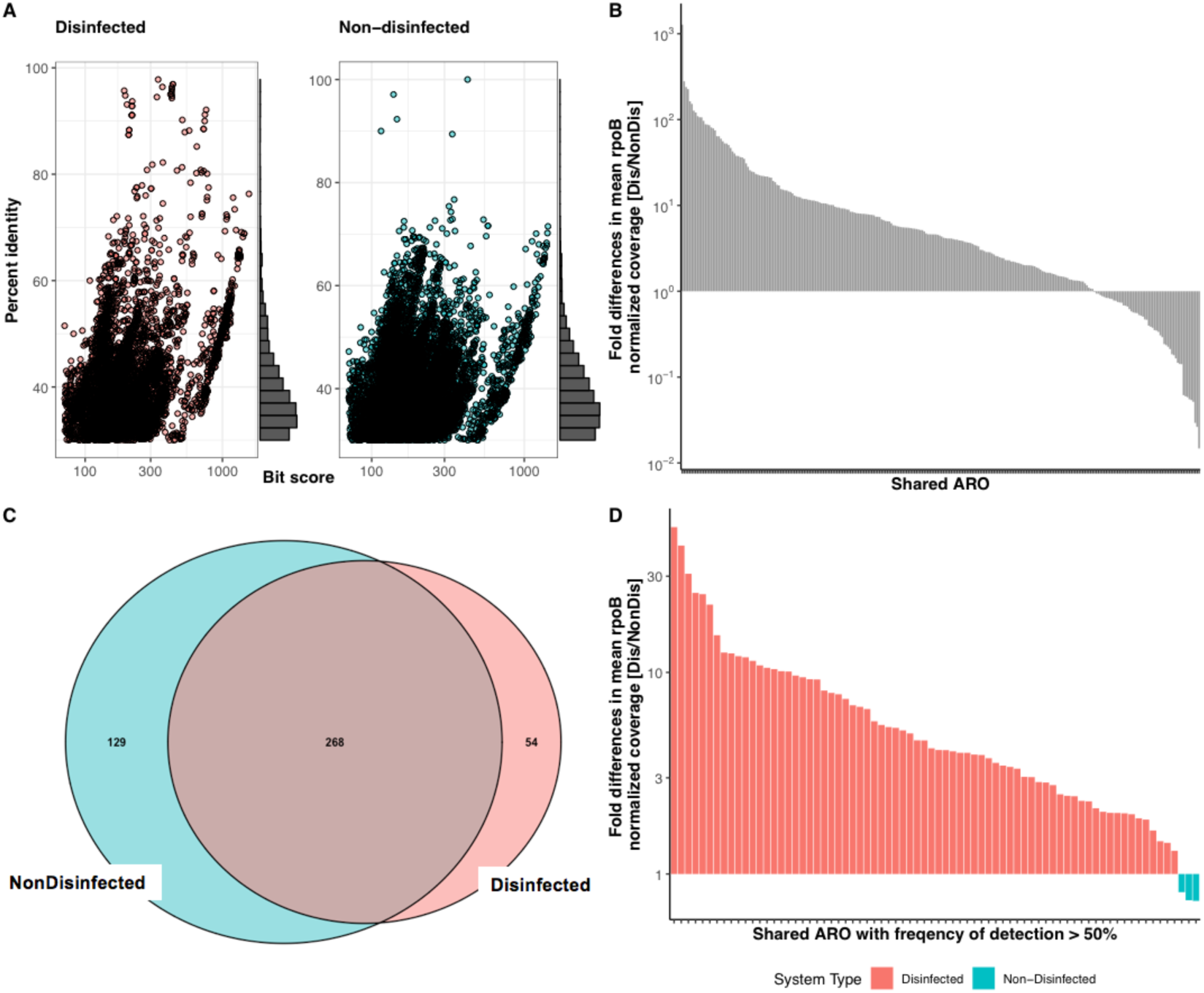
(A) Bitscore and percent identify of CARD annotated ORFs showed similar distributions for Dis (red) and NonDis samples (blue). (B) Fold difference in mean *rpoB* normalized AROs shared between Dis and NonDis systems indicated shared AROs were enriched in Dis systems. (D) Venn diagram of CARD annotated AROs in Dis and NonDis systems indicated significant shared fraction with fewer unique AROs in Dis systems. (D) Fold difference in mean *rpoB* coverage of shared AROs with frequency of detection > 50% also indicated enrichment in Dis samples. Color of bars denotes contribution from disinfection system (Red: Dis and Blue: NonDis).

### Detection frequency of ARO was associated with its relative abundance in disinfected and non-disinfected systems

There was a positive relationship between the *rpoB* normalized coverage and detection frequency of AROs in both Dis and NonDis systems (Figure 2A), with 9 and 37 AROs detected in all Dis and NonDis samples, respectively. Of the total 322 AROs detected in Dis systems, 92 AROs were present in 50% or more samples, whereas 136 out of 397 AROs that were detected in NonDis systems had a frequency of detection greater or equal to 50% (Figure 2B). Among the exclusive AROs, only two and six AROs exhibited detection frequency of 50% or greater for the Dis and NonDis samples, respectively. These exclusive and frequent AROs included a fluoroquinolone resistance gene *mfpA* (*qnr* homolog) with a frequency of detection of 57% and a mean *rpoB* normalized coverage of 0.07±0.16 and LRA-5 *beta-lactamase* with frequency of detection of 52% and a mean of 0.034±0.044 in the Dis samples. The six exclusive and frequent AROs in NonDis samples included (1) three dihydrofolate reductase genes (*dfrK, dfrA22*, and *dfrD*) with detection frequencies between 50-55% and mean *rpoB* normalized coverage of 4.3±5×10^−3^, 2.6±3.2×10^−3^, and 1.7±1.8×10^−3^, respectively, (2) two vancomycin resistance genes (*vanB* and *vanRC*) with detection frequencies of 78% and 50% and mean *rpoB* normalized coverage of 4.5±3.5×10^−3^ and 2.3±2.6×10^−3^, respectively and (3) *ermD* conferring the MLSb resistance phenotype with detection frequency of 50% and a mean *rpoB* normalized coverage of 3.6±7.0×10^−3^.

**Figure 2:**
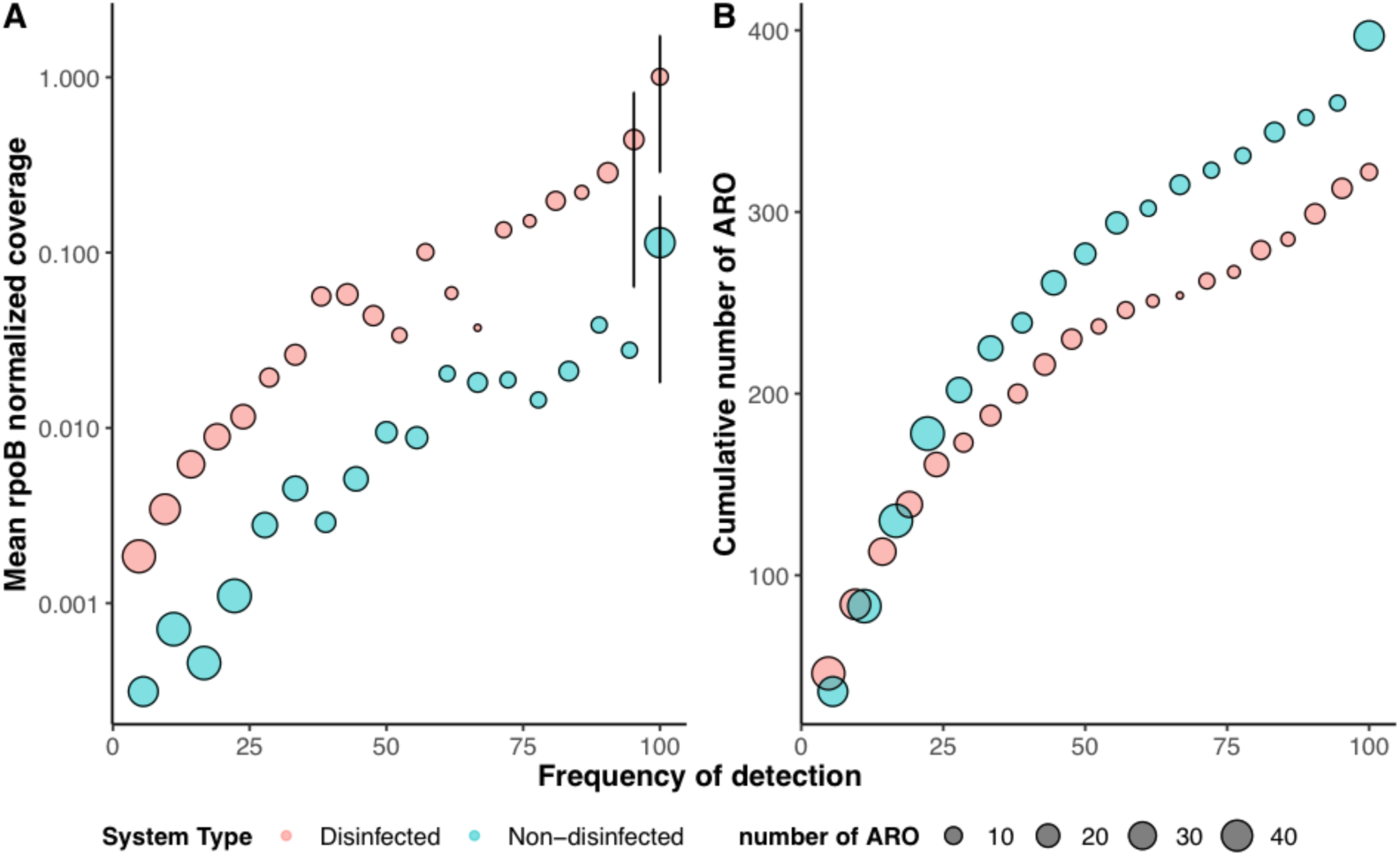
(A) Mean *rpoB* normalized coverage and (B) cumulative number of AROs were positively associated with the frequency of detection of AROs. Red and blue colors correspond to Dis and NonDis samples, respectively and the size of the points represents number of AROs. The mean *rpoB* normalized coverage of AROs at all detection frequencies is higher for Dis systems, although the cumulative number of AROs is generally higher for NonDis systems.

### The presence or absence of disinfectant residual was key in differentiating between resistance profiles of Dis and NonDis systems

The resistance profiles for both Dis and NonDis samples were diverse and even, and showed no significant differences between them (Figure 3A). For instance, while the mean observed AROs were 120 and 160 for the Dis and NonDis samples; the higher mean observed in NonDis samples were due to samples from a single drinking water system (i.e., ND5). This difference can be attributed to greater sequencing depth and higher number of scaffolds post-assembly, and thus higher likelihood of capturing more AROs. To incorporate normalized abundances (i.e., *rpoB* normalized coverage) of the AROs, Simpson and Shannon diversity indexes were calculated converging on the same conclusion.

**Figure 3:**
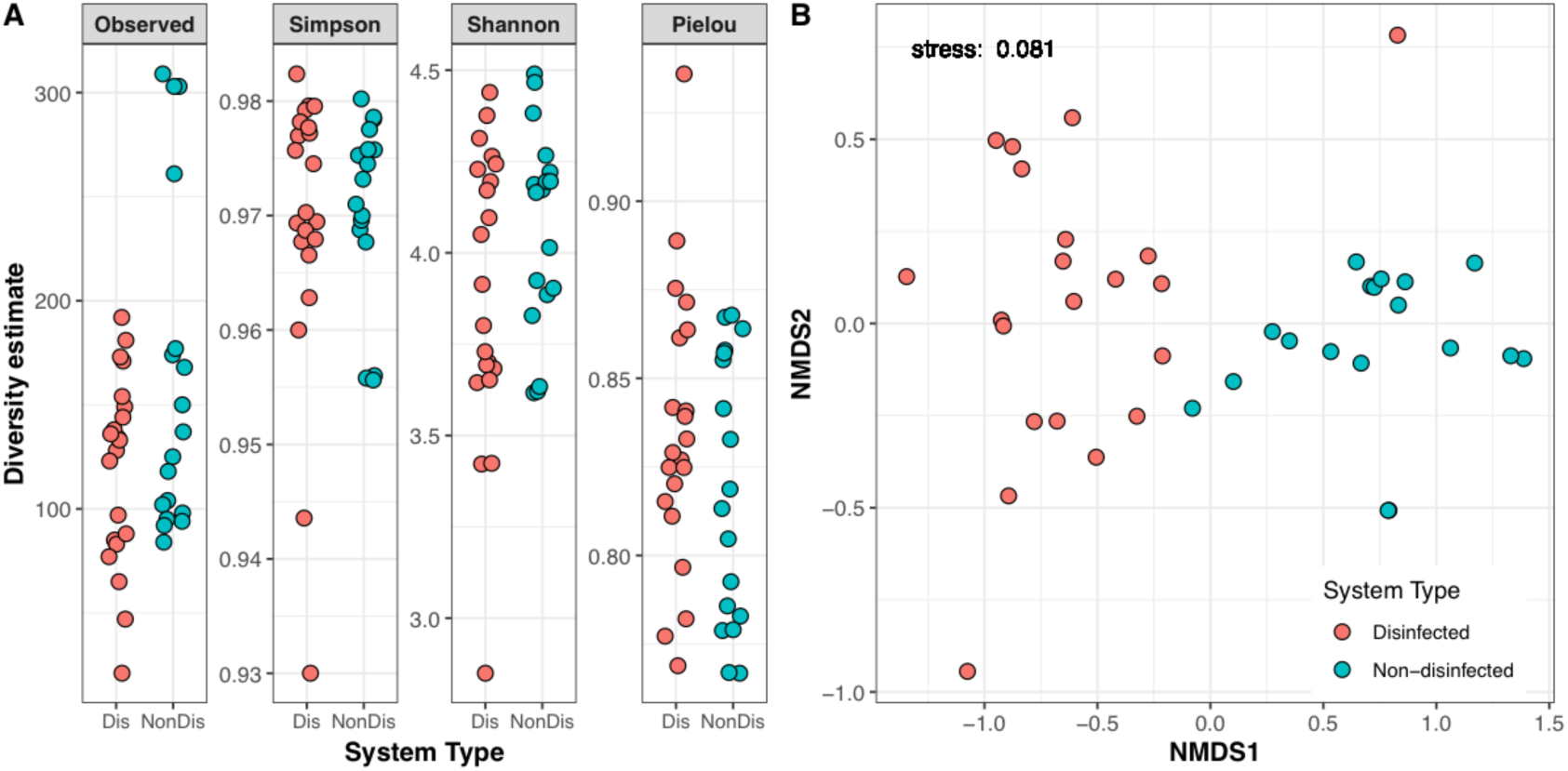
(A) Alpha diversity estimates of samples by strategy. On average, there are 120 AROs observed in Dis systems, and 160 in NonDis systems. Simpson and Shannon indexes indicate highly diverse resistomes, and high Pielou’s evenness index indicates an uneven resistome, although no significant differences are observed for these metrics between strategies. (B) Ordination plot using Bray-Curtis distance describing AROs’ rpoB normalized coverage dissimilarity colored by disinfection strategy (Red: Dis and Blue: NonDis). The clustering suggests that the resistome composition of Dis and NonDis systems is different.

Bray-Curtis distance clustering of samples based on *rpoB* normalized coverage of scaffolds with ARO in each sample showed that the structure of the resistance traits clustered based on the presence/absence of a disinfectant residual (Figure 3B). Permutational ANOVA analyses indicated a significant difference in ARO composition based on grouping of samples based on presence/absence of disinfectant residual (p<0.001). Inclusion of environmental (i.e., temperature) and water chemistry (i.e., conductivity, dissolved oxygen, TOC, nitrate, ammonia, nitrite) along with presence/absence of disinfectant in PERMANOVA explained approximately 57% of the variance suggesting that presence/absence of disinfectant residual is indeed significant (p<0.001) in differentiating ARO composition, while 43% of variability in ARO composition was unexplained (i.e., residuals). However, excluding the disinfectant presence/absence category from PERMANOVA analyses increased the residuals by only 3% with 54% of the variance explained purely based on water chemistry parameters other than the presence/absence of disinfectant residual. This suggests that while the presence/absence of disinfectant residual is a significant factor, the combined effect of water chemistry is also critically important. In contrast to PERMANOVA, post-hoc SIMPER analysis indicated that the overall dissimilarity explained by presence/absence of disinfectant residuals was 54%. The AROs driving these differences were primarily efflux associated genes including *arlR, sav1866, mtrA*, and *golS* and one target alteration gene, *bacA*. These 5 AROs explain 13% of the variation between Dis and NonDis samples, with at least 34 AROs required to explain 50% of the variation between the two sample types.

### Efflux and target alteration are dominant antibiotic resistance mechanisms in disinfected and non-disinfected systems

The AROs detected in this study were associated with six different resistance mechanisms which account for most of the mechanisms represented in the CARD. These include (1) efflux, (2) target alteration, (3) enzymatic inactivation, (4) target protection, (5) target replacement, and (6) reduced permeability (Figure 4A, 4B).

**Figure 4:**
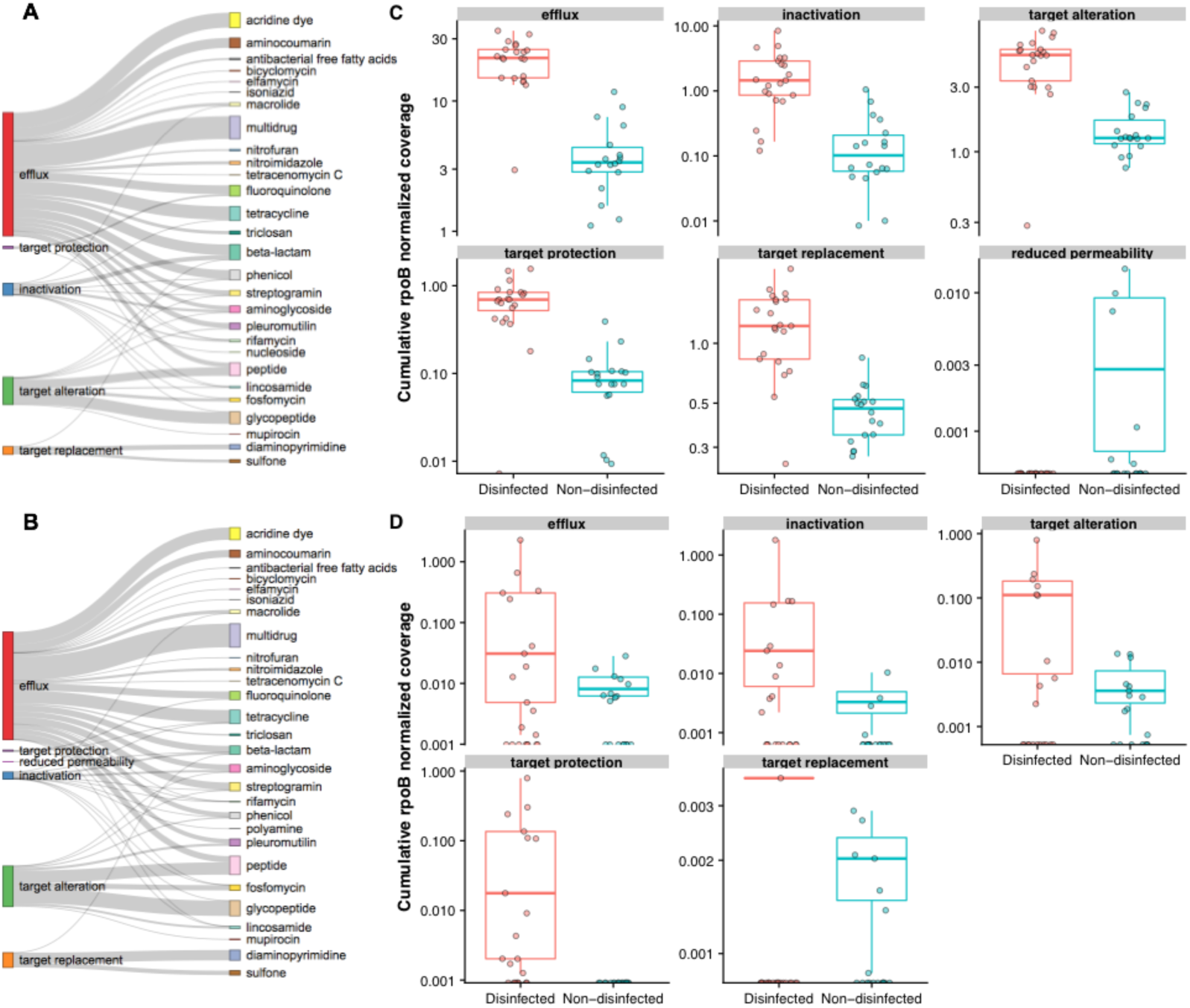
Sankey plot of annotated resistance mechanisms with their corresponding drug class determinants for (A) Dis and (B) NonDis samples. The richness of AROs within mechanisms of resistance exhibits a similar distribution for both systems, with efflux and target alteration mechanisms dominating irrespective of strategy. Comparison of cumulative *rpoB* normalized coverage of mechanisms of resistance between Dis and NonDis samples at (C) “loose” sequence similarity threshold and (D) “stringent” sequence similarity threshold.

Efflux related genes were the most abundant group in both Dis and NonDis samples, 55% and 49% of the AROs in Dis and NonDis samples were categorized as efflux at the “loose” sequence similarity threshold (Figure 4A, 4B). This high prevalence can in large part be attributed to the breadth of functions and diversity of genes that are involved in this mechanism of resistance. The detected efflux genes include those associated with ATP binding cassette (ABC), resistance nodulation cell division (RND), small multidrug resistance (SMR), and major facilitator super family antibiotic efflux (MFS) gene families such as *vga, mex, abe*, and *tet* genes, respectively. Efflux pumps are widespread and present in all organisms^62^, therefore their prevalence and abundance in databases is likely high, as well as the potential for them to be annotated.

Genes associated with target alteration constituted 20.5% and 27.5% of all AROs for Dis and NonDis systems, respectively. Genes in this mechanism category were largely characterized by *erm* (ribosomal alteration) and *van* (cell wall target alteration) genes. Glycopeptide resistance operons (*van* gene clusters) encode enzymes for low affinity precursors of peptidoglycan, and therefore a greater number of individual genes are required if resistance is to be expressed^63^. For example, *vanA* gene cluster contains genes for a two-component regulatory system (*vanR* and *vanS*), three resistance genes (vanH (dehydrogenase, *vanA* (ligase), and *vanX* (DD-dipeptidase)), an accessory gene *vanY* and the *vanZ* gene^64^. Genes involved in enzymatic inactivation of antibiotics, (i.e., acetyltransferases, beta-lactamases, phosphotransferases) accounted for 17.7% and 15.4% of the AROs found in Dis and NonDis samples, respectively with diverse *AAC, Bla*_*OXA*_, and *APH* associated genes.

Genes involved in target protection were less diverse and included almost exclusively *qnr* genes. This mechanism accounted for 1.83% and 1.1% of AROs detected in Dis and NonDis samples, respectively, while they were only detected in Dis samples at the “stringent” sequence similarity threshold. In experiments involving disinfection of wastewater treatment plant (WWTP) effluent followed by regrowth, Di Cesare *et al.* observed that the disinfection process influenced the regrowth of organisms carrying *qnrS* gene. Specifically, chlorinated samples showed increasing growth post disinfection, whereas microbial abundance remained stable in samples from WWTPs using peracetic acid and UV disinfection, respectively^65^.

Target replacement mechanism is more prevalent in NonDis samples at “stringent” sequence similarity threshold. Specifically, genes associated with target replacement constituted 5.0% and 6.5% of the AROs in the Dis and NonDis samples, respectively, at the “loose” sequence similarity threshold, while this changed to 1.1% and 20.0% of the AROs found in the Dis and NonDis samples, respectively, at the “stringent” sequence similarity threshold. The majority of genes associated with this mechanism are *dfr* and *sul* genes which are both related to folic metabolism and are widespread in the environment, being predominantly located on plasmids or transposons^66^. It has been observed that the inactivation of ARGs such as *sul1* from municipal wastewater is most effective with chlorination, followed by UV and ozonation^67^ which could explain the infrequency of detection of these genes in Dis systems. Finally, genes associated with reduced permeability mechanism were only observed under the “loose” sequence similarity threshold for the NonDis samples and only constituted 0.21% of all detected AROs.

While AROs corresponding to each mechanism were present in both, Dis and NonDis sample types, the differential abundance of the mechanisms between the two groups depended on the sequence similarity threshold used for identifying an ARO. Specifically, for all “loose” sequence similarity threshold, all mechanisms were significantly different (p<0.01) between Dis and NonDis samples (Figure 4C), while at the “stringent” sequence similarity threshold these differences were statistically significant only for target protection mechanism (p <0.0001) (Figure 4D).

### Significantly different drug classes are categorized as clinically important and indicate an association with mycobacteria in Dis systems

Mechanisms of resistance are important to ascertain pathways as target for drug development and in fact disinfectants often share mechanisms of action with antibiotics^68^. However, drug classes ultimately dictate clinical treatment and actionable response. The aforementioned mechanisms of resistance were associated with twenty-nine drug classes (Figure 4A, 4B). AROs related to beta-lactams (n=78) and aminoglycosides (n=59) were more diverse and largely associated with antibiotic inactivation as their main mechanism of resistance (Figure 5A, 5B). This is perhaps due to the fact that the CARD has a relatively high contribution of beta-lactam and aminoglycoside resistance sequences in the database^31^, thus they’re more likely to be identified. However, the distribution of annotated drug classes and mechanisms from Dis and NonDis systems in our dataset (Figure 4A, 4B) differs from the CARD distribution. Resistance traits associated with phenicol (33 out of 451 AROs, 33/451), multidrug (28/451), and tetracycline (38/451), as well as streptogramin (29/451) and glycopeptide (45/451) drug classes were also diverse and correspond largely to efflux (phenicol: 25/181, multidrug: 28/181, and tetracycline: 36/181), and target alteration mechanisms (streptogramin: 21/94 and glycopeptide: 44/94), respectively. Resistance traits associated with “other” drug classes (i.e., antibacterial free fatty acids, biciclomycin, elfamycin, mupirocin, nitrofuran, nitroimidazole, nucleoside, polyamine, and tetracenomycin C) were diverse, however when aggregated they were more prevalent and abundant in Dis samples and were associated primarily with efflux mechanisms. For nearly all drug classes, the resistance traits were more diverse and abundant in Dis samples as compared to NonDis samples.

**Figure 5:**
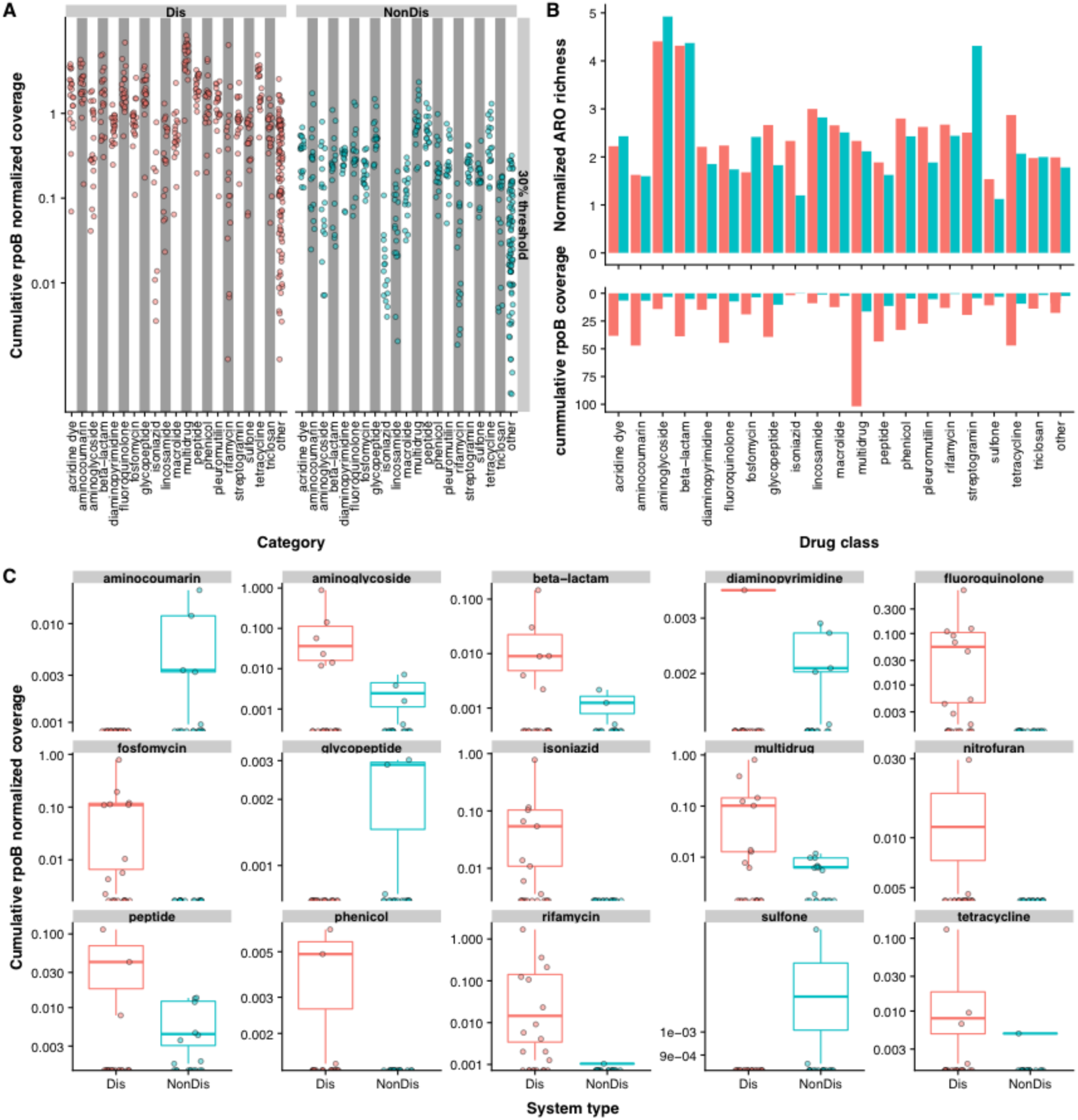
(A) Cumulative *rpoB* normalized coverage of samples within drug classes at “loose” sequence similarity threshold in Dis and NonDis systems. (B) ARO richness normalized by number of samples per strategy for corresponding drug class (top) and cumulative *rpoB* normalized coverage of drug classes in all Dis and NonDis samples (bottom). (C) Comparison of cumulative *rpoB* normalized coverage of drug classes present at “stringent” sequence similarity threshold per disinfection strategy. Colors correspond to disinfection strategy (Red: Dis and Blue: NonDis).

Groupwise comparisons of cumulative mean *rpoB* normalized coverage in drug classes for “stringent” sequence similarity threshold indicated statistically significant differences (p<0.01) between Dis and NonDis groups for resistance traits associated with aminocoumarin, isoniazid, fluoroquinolone, fosfomycin, and rifamycin resistance (Figure 5C). The World Health Organization (WHO) has classified quinolones, fosfomycins and rifamycin (ansamycins) as critically important antimicrobials used in human medicine^69^. Aminocoumarins inhibit DNA gyrase associated with cell division and aminocoumarin producing microbes, such as *Streptomyces spp.*, exhibit intrinsic resistance to it. Within our dataset all genes associated with resistance to this drug class were part of efflux complexes (e.g. *mdt, mux*). At “stringent” sequence similarity threshold, this drug class was only present in NonDis systems, however at “medium” sequence similarity threshold, this drug class is present in both systems. Taxonomic annotation of scaffolds containing AROs associated with this aminocoumarin resistance indicated that majority of them (17 out of 21) originate from *Streptomyces spp*. in NonDis systems

AROs associated with isoniazid, fluoroquinolones, fosfomycin, and rifamycin were significantly more abundant in Dis as compared to NonDis samples (p<0.0001) (Figure 5C). Isoniazid is a prodrug that inhibits cell wall synthesis (i.e. mycolic acids) in *Mycobacterium spp.* It is often used in combination with rifampicin (i.e., rifamycin derivative) to potentiate bactericidal effects. Resistance traits associated with isoniazid resistance were exclusive to Dis systems at both “stringent” and “medium” sequence similarity threshold and mapped back to a single resistance gene *efpA*, which is an efflux pump found in mycobacterial species^70^. Fluoroquinolones interfere with DNA replication and transcription as their primary antimicrobial activity mechanisms. They are synthetic quinolone antibiotics (whose precursor is nalidixic acid) that are effective against Gram-positive and Gram-negative bacteria, generally targeting topoisomerase IV and DNA gyrase, respectively^71^. In this study, mechanisms that mediate resistance to this drug were primarily associated with efflux and target protection, with *emr, pat*, and *qep*, and *qnr* genes as examples, respectively. AROs related to fluoroquinolones were not present in NonDis samples, but were observed in 48% of the samples for Dis samples at “stringent” sequence similarity threshold. These AROs were, however, present in Dis and NonDis samples at “medium” (Dis: 91% of samples; NonDis: 83% of samples) and “loose” (Dis: 100% of samples; NonDis: 100% of samples) sequence similarity threshold (Figure 5A, 5B). Fosfomycin is a broad-spectrum bactericidal drug that inhibits bacterial cell wall biogenesis by inactivating the enzyme UDP-N acetylglucosamine-3-enolpyruvyltransferase, also known as MurA^72^, with *Streptomyces spp.* being the known producers of this antibiotic. Our dataset contains diverse Fos resistance genes that mediate resistance through antibiotic inactivation, and *murA* genes that provide intrinsic resistance through target alteration. Resistance to this drug class was unique to Dis systems at “stringent” sequence similarity threshold. Rifamycin’s mode of action involved inhibition of RNA synthesis and this drug can be biosynthesized by *Amycolaptosis* spp. via a *rif* cluster. This drug class is particularly effective against *Mycobacterium spp.* and some *Mycobacterium spp.*, and are known to have mutations that confer resistance to rifampicin (a rifamycin derivative). In our study, *efr* and *arr* antibiotic resistance genes that mediate resistance via efflux and inactivation mechanisms to rifamycin, respectively, were exclusive to Dis systems at “stringent” sequence similarity threshold. Resistance drug classes that were statistically different between Dis and NonDis systems at the “stringent” sequence similarity threshold were overwhelmingly associated with antibiotics used for mycobacterial infections and more so nearly exclusive to Dis systems. For example, isoniazid is exclusively used for treatment of mycobacterial infections, fluoroquinolone’s precursor nalidixic acid is used as supplement to media supporting mycobacterial growth, fosfomycin resistance *murA* gene confers intrinsic resistance to *Mycobacterium spp.*, and rifamycin *arr* resistance gene also provides intrinsic resistance to *Mycobacterium spp*.

### Distinct host association of AROs in Dis and NonDis systems

Only 16 ARO containing scaffolds were annotated as being of viral origin using either of the approaches that rely on the IMG/VR database, while no ARO containing scaffolds were annotated as viral scaffolds using NCBI or VirFinder. Three out of the 16 AROs containing scaffolds annotated as viral scaffolds contained two AROs each; i.e. *dfrA3* and *vanHD*, MexF and *oqxA*, and *rosB* and *mtrA*, respectively, while the remaining contained a single ARO. The percent identity threshold of AROs on these scaffolds to the CARD was consistently in the “loose” sequence similarity threshold and we describe the results obtained within this context. While our sampling protocol did not address viral recovery, our results suggest that particle (size ≥0.22 µm) and host-associated mobile genetic elements (i.e., viruses and plasmids) do not contribute significantly to AMR traits in drinking water systems. A BLAST alignment of all scaffolds containing AROs (Dis=5,445 NonDis= 9,719) was performed against a local database of bacterial genomes (see Methods section). The alignment of these “loose” sequence similarity threshold scaffolds resulted in only 265 matches at 90% identity and 90% query coverage criteria. Of these, 263 scaffolds were from Dis systems and matched with sequences belonging to the genera *Mycobacterium* (n=254), *Pseudomonas* (n=5), *Legionella* (n=3), *Burkholderia* (n=1), *Aeromonas* (n=1), and *Salmonell*a (n=1). Only two scaffolds from the NonDis systems matched reference genomes within the genera *Pseudomonas* and *Salmonella* (Figure 6A). A subsequent web-based BLAST alignment was consistent with the BLAST results against the local database.

**Figure 6:**
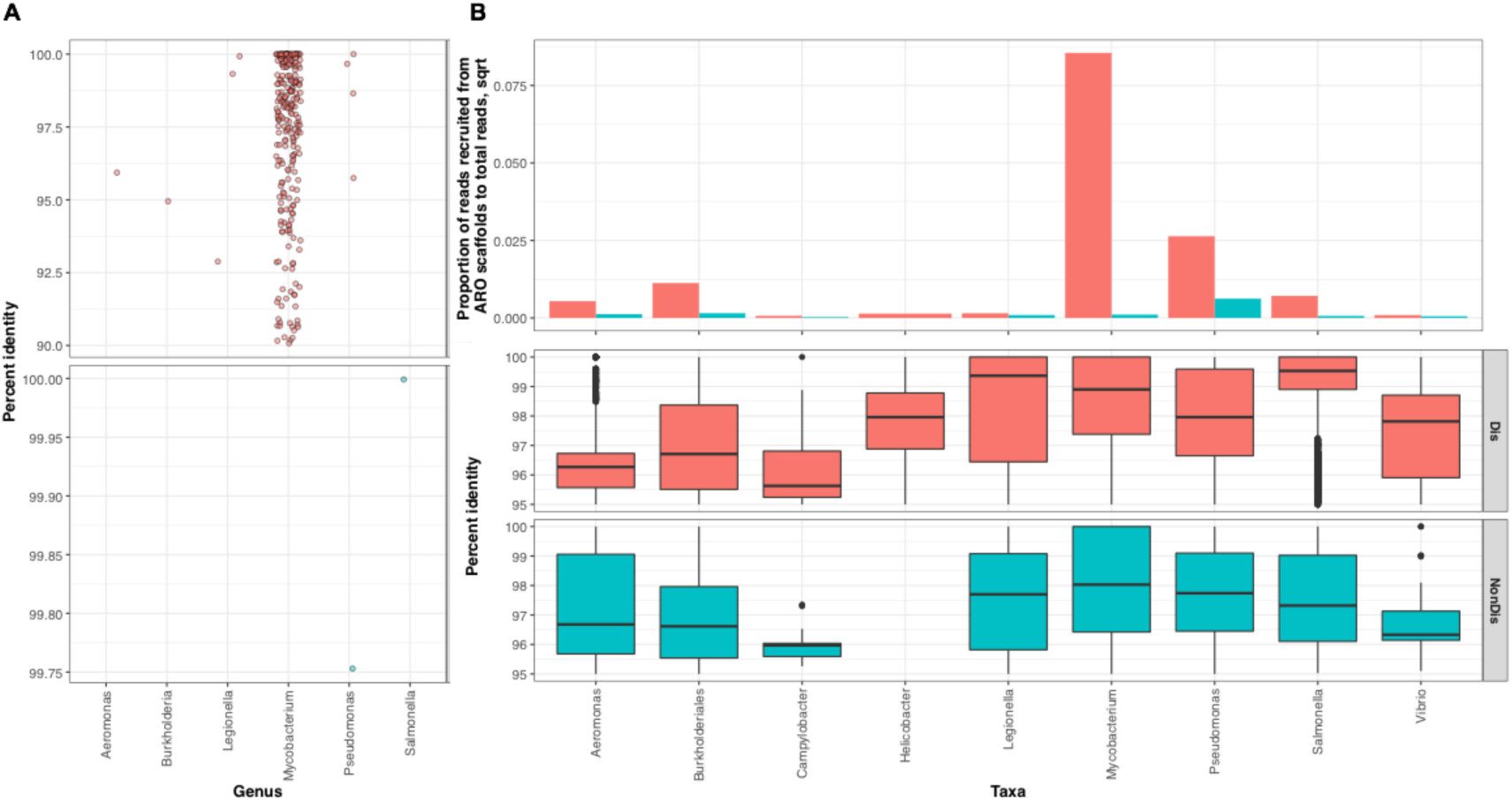
(A) Scaffold-based BLAST annotation to local pathogen database for ARO carrying scaffolds showed that the majority of the annotated scaffolds correspond to Dis systems and *Mycobacterium spp*. (B) Read based BLAST annotation of unclassified scaffolds from (A) with corresponding total reads per disinfection strategy. A distinct host association of AROs was observed for Dis and NonDis systems, where *Mycobacterium spp.* are predominant in Dis systems and *Pseudomonas spp*. in NonDis.

The majority of the scaffolds with loose identity threshold AROs (n=14,899) did not match any of the reference genomes at the aforementioned criteria. To further inspect the taxonomic identity of these scaffolds, we extracted reads mapping to these scaffolds (n=134,955,066, Dis=117,798,239 and NonDis = 17,156,827) and aligned them against the reference pathogen genome database using stringent criteria of 95% sequence similarity with a 95% query coverage using BLAST. Approximately 1.5% of the extracted reads mapped to the reference genomes with a majority of these associated with scaffolds from Dis systems (n=2,050,992) as compared to the NonDis systems (n=10,484). A total of 88.3% of the reads extracted from scaffolds containing AROs from Dis systems mapped to mycobacterial genomes in the reference database. However, these reads constituted 0.72% of all reads from Dis systems. In contrast, 84.7% of the reads extracted from ARO containing scaffolds from NonDis systems mapped to *Pseudomonas* genomes, which constituted 0.004% of all reads from NonDis systems (Figure 6B). This suggests that the ARO association with known genomes in potential pathogenic genera was significantly lower in NonDis systems compared to Dis systems, with mycobacteria being the primary host of antibiotic resistance traits in Dis systems. However, while AMR containing mycobacteria were enriched in Dis systems, this does not equate to higher quantitative exposures as the typical absolute abundance (i.e., concentration) of microbes in Dis systems is two to three orders of magnitude lower than that in NonDis systems. Nevertheless, the results agree to some extent with previous findings that suggest low sequence similarity between *hsp65* genes found in NonDis systems in the Netherlands and *hsp65* from opportunistic mycobacteria^73^, as *Mycobacterium spp.* constituted 2.55% of the reads extracted from scaffolds containing AROs and pertaining to NonDis systems which represented < 0.0001% of all reads pertaining to NonDis systems in this study.

### Genome-resolved metagenomics indicates that ARO harboring mycobacteria are responsive to chlorine concentrations

To further understand the ARO association with mycobacteria in Dis systems, we recovered five mycobacterial MAGs from the D3, D4, and D5 metagenome assemblies. The complete statistics associated with each mycobacterial MAG are presented in Table 1. Genome taxonomy database annotation indicated that Bins 4 and 13 were closely related to *Mycobacterium llatzerense* with 99.4 and 97.0% sequence average nucleotide identity to the same draft genome (GCF_000746215) which was assembled from plant associated samples. Bin 3 exhibited 99.4% sequence similarity to a mycobacterial isolate sampled from tap water sample in Germany (GCF_002013415), while Bins 9 and 10 did not have any closely related mycobacterial genomes at the species level. Phylogenetic placement of the bins based on both *rpoB* and multigene alignment of ribosomal proteins indicated that all assembled bins belonged to environmental and rapidly growing mycobacteria (Figure 7). While Bin 3 placed in the rapid growing *Mycobacterium chelonae*-*absceccus* complex, Bins 4 and 13 were closely phylogenetically clustered with *M. llatzerense*. Although Bins 9 and 10 were distantly related from most mycobacterial reference genomes, they were closely associated with mycobacterial isolates from hydrocarbon contaminated and river estuary sediment, respectively. All the five assembled MAGs contained at least one ARO, with Bins 10 and 13 containing three and two, respectively (Table 1). Specifically, the identified AROs were RbpA, *mtrA*, and *murA*. In two of these bins, RbpA (rifamycin resistance) and *mtrA* (macrolife and penam resistance) co-occur. *M. llatzarense* has been isolated from DWDS and is described as a NTM that resists amoebal phagocytosis and survives disinfection^74,75^. Similarly, Bin3 associated with the *M. chelonae*-*absceccus* complex also demonstrated the presence of gene conferring rifamycin resistance. The enrichment of mycobacterial MAGs with AMR traits in Dis systems can be attributed to several factors. For instance, mycobacteria are known to harbour intrinsic antibiotic resistance^76^ and their colonization, survival, and persistence in DWDS and their resistance to disinfectants at low nutrient concentrations has been well documented^77–79^. Further, intracellular colonization of mycobacteria in protozoa may allow for survival and proliferation of these genera^80^ and sheltering within biofilms can also enhance their survival^79,81–83^.

**Table 1:** Statistics associated with recovered bins along with taxonomy and resistance profile information.

| MAG | Assembly | Completeness (%) | Redundancy (%) | Size (Mbp) | Closest reference genome |  |  | Resistance gene | AMR Gene family | Mechanism | Drug class |
| --- | --- | --- | --- | --- | --- | --- | --- | --- | --- | --- | --- |
|  |  |  |  |  |  | ANI | AF |  |  |  |  |
| Bin3 | D3 | 69.57 | 3.56 | 4.07 | *Mycobacterium spp.* (GCF_002013415.1) | 99.36 | 0.84 | RbpA | RNA polymerase binding protein | Target protection | Rifamycin |
| Bin4 | D5 | 63.07 | 3.99 | 5.58 | *Mycobacterium llatzerense* (GCF_000746215.1) | 96.25 | 0.7 | mtrA | Resistance-nodulation-cell division | Efflux | Macrolide, penam |
| Bin9 | D3 | 71.45 | 1.16 | 4.18 | *Mycobacterium spp.* (N/A) | N/A | N/A | mtrA | Resistance-nodulation-cell division | Efflux | Macrolide, penam |
| Bin10 | D4 | 98.49 | 0.00 | 5.76 | *Mycobacterium spp.* (N/A) | N/A | N/A | RbpA | RNA polymerase binding protein | Target protection | Rifamycin |
|  |  |  |  |  |  |  |  | murA | murA transferase | Target alteration | Fosfomycin |
|  |  |  |  |  |  |  |  | mtrA | Resistance-nodulation-cell division | Efflux | Macrolide, penam |
| Bin13 | D4 | 90.02 | 0.97 | 4.86 | *Mycobacterium llatzerense* (GCF_000746215.1) | 97.02 | 0.89 | mtrA | Resistance-nodulation-cell division | Efflux | Macrolide, penam |
|  |  |  |  |  |  |  |  | RbpA | RNA polymerase binding protein | Target protection | Rifamycin |

**Figure 7:**
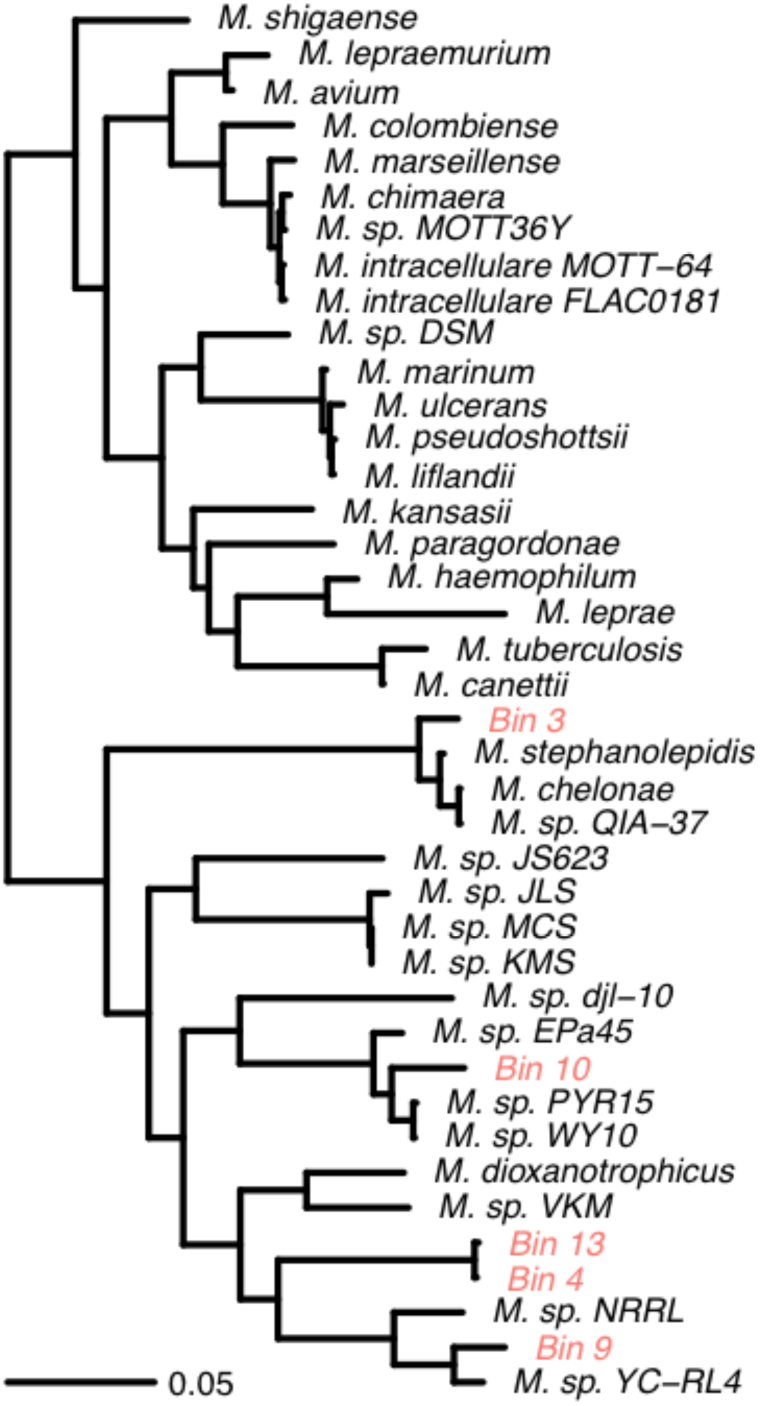
Phylogenomic tree contextualizing selected MAGs (in red font) and selected *Mycobacterium* spp. generated in anvi’o by extracting 48 single-copy ribosomal protein genes from MAGs and 35 complete mycobacterial reference genomes (downloaded from NCBI). A multi gene alignment was generated for each MAG or reference genome from extracted ribosomal protein genes and it was followed by phylogenetic tree reconstruction using FastTree.

In conclusion, in this study we have identified and characterized the AMR traits of tap water from drinking water distribution systems (DWDS) with and without disinfectant residual, Dis and NonDis, respectively. We observed that presence/absence of disinfectant plays a significant role in AMR trait prevalence, composition, and abundance. Although both systems have a diverse and uneven AMR trait distribution, they were more abundant in Dis systems compared to NonDis systems. ARO drug classes that are significantly different between Dis and NonDis systems are associated with scaffolds classified as mycobacteria that were detected exclusively in Dis systems. The overall findings of our study indicate that AMR harboring NTM are systematically enriched in Dis systems. An important direction for future research on is coupling qPCR of ARGs in DWDS with functional metagenomics to determine absolute concentrations of ARGs and assess if the associated hosts are active and AMR traits are functional. This information will be vital to translate our characterization of the AMR traits in drinking water systems to quantitative microbiological risk assessment.

## Author contributions

MS: sample collection and processing and data analyses; ZD: data analyses; SC: sample processing; QMBS: sample collection and processing; AME: supported data analyses, PW: sample collection support, UZI: provided bioinformatic support, AJP: Conceived and designed the analyses, collected data, data analyses. All authors contributed to data interpretation and manuscript writing.

## Conflict of interest Statement

The authors declare no competing financial interests

## Data availability

DNA-sequencing data are available from SRA under BioProject ID PRJNA533545

## Acknowledgements

MS was supported by the College of Engineering at Northeastern University. This work was funded by Engineering and Physical Science Research Council (EP/M016811/1) and the National Science Foundation (NSF-CBET 1749530). UZI is supported by NERC Independent Research Fellowship (NERC NE/L011956/1)

