## Supplementary_text for "Disinfectant residuals in drinking water systems select for mycobacterial populations with intrinsic antimicrobial resistance"

**Supplementary text 1: Identification and removal of contaminant contigs from metagenome assemblies**. Reads obtained from negative controls with spiked mock communities were mapped to genomes of mock community members using bwa-mem and unmapped reads were extracted as contaminant reads using samtools view (flag –f4). Subsequently, reads from each sample and contaminant reads from negative controls were mapped to scaffolds from each metagenomic assembly. The coverage of each scaffold in the sample and the negative control was calculated using genomcov function in bedtools. The relative abundance (RA) of each scaffold was calculated as the coverage of each scaffold divided by the sum of coverage of all scaffolds in the sample, including coverage of spiked mock community organisms for the negative control. Relative standard deviation (RSD) of the coverage across each scaffold in each sample and negative controls was determined by dividing the standard deviation in coverage across the length of each scaffold by its mean coverage in the sample/negative control. The RSD metric was used as a measure of stability of coverage of a scaffold in a sample. A scaffold was considered present in a sample if it was not detected in the negative controls (i.e., negative control coverage = 0) or if its RA_sample_ > RA_control_ and RSD_sample_ < RSD_control_. Any scaffold that did not meet either of these criteria was considered as a contaminant scaffold and was removed from any further sample analyses.

**Supplementary text 2: Construction of potential waterborne pathogen database**. A string search of names included in a WHO list of waterborne pathogens relevant to water supplies (<https://www.who.int/water_sanitation_health/gdwqrevision/watpathogens.pdf>) was used to download fna files excluding “virus” and “phage” string matches from NCBI’s refseq (<ftp://ftp.ncbi.nlm.nih.gov/genomes/ASSEMBLY_REPORTS/assembly_summary_refseq.txt>, last accessed Jul 14 2018). A total of 50312 genbank files (fna.gz) were recruited from Acanthamoeba (24), Acinetobacer (3711), Aeromonas (329), Burkholderia (2272), Campylobacter (2553), Cryptosporidium (39), Cyclospora (18), Dracunculus (1), Entamoeba (10), Eschericia (12598), Giardia (8), Legionella (715), Mycobacterium (7148), Naegleri (2), Pseudmonas (5517), Salmonella (9242), Schistosoma (7), Shigella (1903), Stenotrophomnas (493), Toxoplasma (17), Vibrio (2879), Yersinia (826) , with the respective contribution of each string shown in parenthesis. The blast database was created for these genomes using makeblastdb command.
