## Supplementary_tables for "Disinfectant residuals in drinking water systems select for mycobacterial populations with intrinsic antimicrobial resistance"

**Table S1**: Details on samples, sampling locations, and water chemistry parameters measured for the collected samples.

| **Sample** | **DWS** | **Country** | **Temp (ºC)** | **pH** | **Condcutivity**  **(mS/cm)** | **DO**  **(mg/l)** | **Total**  **Chlorine (mg Cl_2_/l)** | **Phosphate**  **(mg PO_4_^3-^/l)** | **TOC**  **(mg/l)** | **Ammonia**  **(mg N/l)** | **Nitrate**  **(mg N/l)** | **System**  **Type** |
| --- | --- | --- | --- | --- | --- | --- | --- | --- | --- | --- | --- | --- |
| D1.1 | D1 | UK | 16.6 | 8.54 | 128.9 | 4.99 | 0.36 | 1.09 | 2.29 | 0.02 | 0.60 | Dis |
| D1.2 | D1 | UK | 18.3 | 8.51 | 116.5 | 5.07 | 0.31 | 0.98 | 2.15 | 0.04 | 0.48 | Dis |
| D1.3 | D1 | UK | 16.6 | 8.47 | 113 | 5.41 | 0.10 | 1.12 | 1.94 | 0.03 | 0.42 | Dis |
| D1.4 | D1 | UK | 18.3 | 8.39 | 114.5 | 5.54 | 0.13 | 1.15 | 2.29 | 0.03 | 0.49 | Dis |
| D2.1 | D2 | UK | 13.3 | 8.39 | 65.5 | 10.76 | 0.73 | 2.44 | 2.64 | 0.08 | 0.21 | Dis |
| D2.2 | D2 | UK | 12.5 | 8.45 | 65 | 10.05 | 0.42 | 2.06 | 2.62 | 0.08 | 0.17 | Dis |
| D2.3 | D2 | UK | 12.3 | 8.42 | 66 | 10.1 | 0.43 | 2.10 | 2.72 | 0.08 | 0.37 | Dis |
| D2.4 | D2 | UK | 14.5 | 8.33 | 68.7 | 11.36 | 0.33 | 2.07 | 2.44 | 0.07 | 0.25 | Dis |
| D3.1 | D3 | UK | 19.9 | 7.88 | 144.75 | 10.37 | 0.16 | 1.23 | 1.39 | <0.02 | 1.49 | Dis |
| D3.2 | D3 | UK | 10.6 | 7.54 | 148.25 | 11.9 | 0.26 | 1.82 | 1.91 | <0.02 | 1.53 | Dis |
| D4.1 | D4 | UK | 15.1 | 7.56 | 313 | 8.92 | 0.66 | 1.97 | 1.80 | <0.02 | 1.10 | Dis |
| D4.2 | D4 | UK | 17.4 | 7.67 | 312.5 | 9.46 | 0.50 | 2.00 | 1.60 | <0.02 | 1.01 | Dis |
| D4.3 | D4 | UK | 18.3 | 7.95 | 311.5 | 9.17 | 0.60 | 2.12 | 1.63 | <0.02 | 1.07 | Dis |
| D4.4 | D4 | UK | 17.9 | 7.93 | 306 | 9.59 | 0.36 | 1.96 | 1.58 | <0.02 | 1.00 | Dis |
| D5.1 | D5 | UK | 19.2 | 7.49 | 666 | 8.11 | 0.63 | 4.94 | 2.32 | 0.14 | 6.07 | Dis |
| D5.2 | D5 | UK | 18.2 | 7.54 | 683.5 | 13.26 | 0.42 | 0.85 | 2.24 | <0.02 | 6.02 | Dis |
| D5.3 | D5 | UK | 21.1 | 7.92 | 652 | 8.86 | 0.36 | 4.67 | 3.61 | 0.13 | 5.68 | Dis |
| D5.4 | D5 | UK | 21.6 | 7.47 | 653 | 8.09 | 0.42 | 4.77 | 2.28 | <0.02 | 5.82 | Dis |
| D6.1 | D6 | UK | 17.9 | 7.88 | 216.4 | 9.68 | 0.38 | 3.06 | 1.00 | <0.02 | 1.70 | Dis |
| D6.2 | D6 | UK | 19.4 | 8.08 | 212.6 | 9.13 | 0.12 | 3.05 | 0.94 | <0.02 | 1.64 | Dis |
| D6.3 | D6 | UK | 19.7 | 8.02 | 227.65 | 9.33 | 0.26 | 3.09 | 1.08 | <0.02 | 1.73 | Dis |
| ND1.1 | ND1 | NL | 11.4 | 8.25 | 542 | 8.98 | 0 | 0.03 | 1.01 | <0.02 | 0.62 | NonDis |
| ND1.2 | ND1 | NL | 11.8 | 8.23 | 546 | 8.74 | 0 | 0.03 | 1.01 | - | 0.60 | NonDis |
| ND1.3 | ND1 | NL | 12.0 | 8.24 | 546 | 8.7 | 0 | 0.03 | 1.09 | <0.02 | 0.61 | NonDis |
| ND1.4 | ND1 | NL | 14.8 | - | - | - | 0 | 0.03 | 1.04 | - | 0.55 | NonDis |
| ND2.1 | ND2 | NL | 12.4 | 8.11 | 467 | 8.41 | 0 | <0.03 | 2.80 | <0.02 | 0.99 | NonDis |
| ND2.2 | ND2 | NL | 11.8 | 8.13 | 472 | 8.33 | 0 | 0.03 | 2.72 | <0.02 | 0.91 | NonDis |
| ND2.3 | ND2 | NL | - | - | - | - | 0 | <0.03 | 2.63 | - | 0.93 | NonDis |
| ND3.1 | ND3 | NL | 11.7 | 8.09 | 619 | 8.5 | 0 | <0.03 | 5.60 | <0.02 | 2.78 | NonDis |
| ND3.2 | ND3 | NL | 11.3 | 8.06 | 599 | 8.5 | 0 | <0.03 | 5.50 | <0.02 | 2.84 | NonDis |
| ND3.3 | ND3 | NL | 10.7 | 7.97 | 618 | 9.1 | 0 | <0.03 | 5.40 | <0.02 | 2.76 | NonDis |
| ND3.4 | ND3 | NL | 10.1 | 8.06 | 614 | 8.7 | 0 | <0.03 | 5.40 | <0.02 | 2.98 | NonDis |
| ND4.1 | ND4 | NL | 11.9 | 8.37 | 322 | 10.2 | 0 | <0.03 | 1.00 | <0.02 | 0.48 | NonDis |
| ND4.2 | ND4 | NL | 11.4 | 8.28 | 336 | 10.3 | 0 | <0.03 | 1.10 | <0.02 | 0.53 | NonDis |
| ND4.3 | ND4 | NL | 10.8 | 8.38 | 334 | 9.7 | 0 | <0.03 | 1.00 | <0.02 | 0.48 | NonDis |
| ND5.1 | ND5 | NL | 11.6 | 7.59 | 507 | 8.6 | 0 | <0.03 | 3.90 | <0.02 | 2.59 | NonDis |
| ND5.2 | ND5 | NL | 10.3 | 8.01 | 527 | 9.3 | 0 | <0.03 | 3.80 | <0.02 | 0.96 | NonDis |
| ND5.3 | ND5 | NL | 10.7 | 7.61 | 482 | 8.7 | 0 | <0.03 | 4.20 | <0.02 | 2.79 | NonDis |
| ND5.4 | ND5 | NL | 9.5 | 7.53 | 506 | 7.5 | 0 | <0.03 | 3.20 | <0.02 | 2.80 | NonDis |

**Table S2**: Names and DSM catalogue numbers of bacteria used to construct mock communities and their corresponding accession numbers.

| **Organism name** | **DSM catalog no.** | **Genbank assembly accession no.** |
| --- | --- | --- |
| *Beijerinckia india* | 1715 | CP00101 |
| *Desulfosporosinus orientis* | 765 | CP003108 |
| *Flectobacillus major* | 103 | ATXY00000000 |
| *Legionella pneumophila* | 7513 | AE017354 |
| *Listeria monocytogenes* | 19094 | HE999705.1 |
| *Meiothermus ruber* | 1279 | CP001743 |
| *Neisseria meningitides* | 15464 | AM421808 |
| *Propionibacterium acnes* | 16379 | AE017283 |
| *Pseudomonas aeruginosa* | 1128 | NC_009656 |
| *Spirosoma linguale* | 74 | CP001769 |

**Table S3**: Reference mycobacterial genomes used for phylogenomic placement of mycobacterial bins assembled from drinking water samples and their NCBI assembly numbers.

| **Mycobacterial species** | **Assembly number** |
| --- | --- |
| *M. avium* | GCA_001683455 |
| *M. canettii* | GCA_000253375 |
| *M. chelonae* | GCA_003390495 |
| *M. chimaera* | GCA_001750065 |
| *M. colombiense* | GCA_002105755 |
| *M. dioxanotrophicus* | GCA_002157835 |
| *M. haemophilum* | GCA_000340435 |
| *M. intracellulare FLAC0181* | GCA_002285675 |
| *M. intracellulare MOTT-64* | GCA_000276825 |
| *M. kansasii* | GCA_000157895 |
| *M. leprae* | GCA_003253775 |
| *M. lepraemurium* | GCA_002291465 |
| *M. liflandii* | GCA_000026445 |
| *M. marinum* | GCA_003391395 |
| *M. marseillense* | GCA_002285715 |
| *M. paragordonae* | GCA_003614435 |
| *M. pseudoshottsii* | GCA_003584745 |
| *M. shigaense* | GCA_002356315 |
| *M. sp. djl-10* | GCA_001695755 |
| *M. sp. DSM* | GCA_900292015 |
| *M. sp. EPa45* | GCA_001021385 |
| *M. sp. JLS* | GCA_000016005 |
| *M. sp. JS623* | GCA_000328565 |
| *M. sp. KMS* | GCA_000015405 |
| *M. sp. MCS* | GCA_000014165 |
| *M. sp. MOTT36Y* | GCA_000262165 |
| *M. sp. NRRL* | GCA_001580405 |
| *M. sp. PYR15* | GCA_002335685 |
| *M. sp. QIA-37* | GCA_001611855 |
| *M. sp. VKM* | GCA_000416365 |
| *M. sp. WY10* | GCA_001886515 |
| *M. sp. YC-RL4* | GCA_001644575 |
| *M. stephanolepidis* | GCA_002356335 |
| *M. tuberculosis* | GCA_000706665 |
| *M. ulcerans* | GCA_900638745 |

**Table S4**: Sequencing and assembly statistics for drinking water metagenomes

| **Assembly** | **Sample ID** | **Raw reads** | **QC reads** | **% reads mapping** | **% True Scaffolds** | **Assembly size (Mbp)** | **N50** | **L50** | **Scaffolds > 1000 bp** | **#ORFs** |
| --- | --- | --- | --- | --- | --- | --- | --- | --- | --- | --- |
| D1 | D1.1 | 6.80E+07 | 5.55E+07 | 99.03 | 98.4 | 615.10 | 1131 | 85015 | 107347 | 863401 |
|  | D1.2 | 7.00E+07 | 5.78E+07 |  |  |  |  |  |  |  |
|  | D1.3 | 1.05E+07 | 8.93E+06 |  |  |  |  |  |  |  |
|  | D1.4 | 8.87E+07 | 7.34E+07 |  |  |  |  |  |  |  |
| D2 | D2.1 | 2.14E+06 | 1.50E+06 | 96.25 | 95.2 | 53.03 | 2112 | 3515 | 10078 | 75291 |
|  | D2.2 | 1.64E+07 | 1.34E+07 |  |  |  |  |  |  |  |
|  | D2.3 | 1.74E+07 | 1.41E+07 |  |  |  |  |  |  |  |
|  | D2.4 | 2.23E+07 | 1.79E+07 |  |  |  |  |  |  |  |
| D3 | D3.1 | 1.64E+07 | 1.25E+07 | 91.15 | 99.2 | 249.69 | 1531 | 26834 | 54708 | 374097 |
|  | D3.2 | 7.02E+07 | 4.87E+06 |  |  |  |  |  |  |  |
| D4 | D4.1 | 1.20E+07 | 9.25E+06 | 93.04 | 98.9 | 204.78 | 3300 | 7882 | 35929 | 269598 |
|  | D4.2 | 3.26E+07 | 2.20E+06 |  |  |  |  |  |  |  |
|  | D4.3 | 1.14E+07 | 9.39E+06 |  |  |  |  |  |  |  |
|  | D4.4 | 2.12E+07 | 1.52E+07 |  |  |  |  |  |  |  |
| D5 | D5.1 | 1.26E+07 | 9.42E+06 | 88.73 | 99.1 | 269.12 | 1313 | 32603 | 51694 | 410972 |
|  | D5.2 | 1.14E+07 | 8.79E+06 |  |  |  |  |  |  |  |
|  | D5.3 | 1.07E+07 | 8.31E+06 |  |  |  |  |  |  |  |
|  | D5.4 | 1.35E+07 | 1.02E+07 |  |  |  |  |  |  |  |
| D6 | D6.1 | 4.97E+05 | 3.44E+05 | 95.90 | 98.2 | 57.23 | 1641 | 5201 | 12510 | 86083 |
|  | D6.2 | 1.36E+07 | 1.06E+07 |  |  |  |  |  |  |  |
|  | D6.3 | 8.83E+06 | 6.40E+06 |  |  |  |  |  |  |  |
| ND1 | ND1.1 | 1.30E+07 | 1.01E+07 | 83.83 | 99.3 | 472.02 | 855 | 137550 | 96504 | 851154 |
|  | ND1.2 | 1.31E+07 | 1.02E+07 |  |  |  |  |  |  |  |
|  | ND1.3 | 1.75E+07 | 1.33E+07 |  |  |  |  |  |  |  |
|  | ND1.4 | 1.58E+07 | 1.19E+07 |  |  |  |  |  |  |  |
| ND2 | ND2.1 | 9.35E+06 | 7.63E+06 | 75.03 | 99.3 | 316.18 | 802 | 100192 | 58972 | 571361 |
|  | ND2.2 | 1.01E+07 | 7.88E+06 |  |  |  |  |  |  |  |
|  | ND2.3 | 1.33E+07 | 1.05E+07 |  |  |  |  |  |  |  |
| ND3 | ND3.1 | 1.82E+07 | 1.38E+07 | 81.64 | 99.4 | 562.73 | 775 | 194433 | 100086 | 1034341 |
|  | ND3.2 | 9.92E+06 | 8.29E+06 |  |  |  |  |  |  |  |
|  | ND3.3 | 1.65E+07 | 1.30E+07 |  |  |  |  |  |  |  |
|  | ND3.4 | 1.68E+07 | 1.35E+07 |  |  |  |  |  |  |  |
| ND4 | ND4.1 | 4.28E+06 | 2.26E+06 | 66.73 | 99.4 | 143.22 | 808 | 45681 | 27235 | 261063 |
|  | ND4.2 | 4.83E+06 | 2.45E+06 |  |  |  |  |  |  |  |
|  | ND4.3 | 1.73E+07 | 1.31E+07 |  |  |  |  |  |  |  |
| ND5 | ND5.1 | 4.35E+07 | 3.12E+07 | 93.75 | 99.5 | 1834.75 | 1005 | 415411 | 419672 | 3054579 |
|  | ND5.2 | 4.88E+07 | 3.51E+07 |  |  |  |  |  |  |  |
|  | ND5.3 | 3.82E+07 | 2.78E+07 |  |  |  |  |  |  |  |
|  | ND5.4 | 5.11E+07 | 3.69E+07 |  |  |  |  |  |  |  |
